# Mouse navigation strategies for odor source localization

**DOI:** 10.1101/558643

**Authors:** Annie Liu, Andrew E Papale, James Hengenius, Khusbu Patel, Bard Ermentrout, Nathaniel N Urban

**Affiliations:** Department of Neurobiology, University of Pittsburgh School of Medicine, Pittsburgh, PA; Department of Mathematics, University of Pittsburgh, Pittsburgh, PA; University of Pittsburgh, Pittsburgh, PA; Center for the Neural Basis of Cognition, Pittsburgh, PA

**Author notes:** **CORRESPONDING AUTHOR** Andrew E. Papale. These three authors contributed equally to this manuscript (Annie Liu, Andrew E Papale, James Hengenius). **AUTHOR CONTRIBUTIONS** JH, AL, BE, and NNU designed the behavioral experiments. AEP, JH, BE, and NNU designed the simulations. AL and KP conducted the behavioral experiments. AEP, JH, and AL analyzed the data. AEP and JH wrote the MATLAB code. AEP calculated the statistics. AEP, JH, AL, BE, and NNU wrote the paper.

**Keywords:** Olfaction, navigation, casting, binaral-olfaction, binaral-sniffing, serial-sniffing

## Abstract

Navigating an odor landscape is a critical behavior for the survival of many species, including mice. An ethologically relevant mouse behavior is locating food using information about odor concentration, such as the relatively laminar gradient established by diffusion-convection near surfaces. To approximate this behavior, we use an open field odor-based spot-finding task indoors with little wind, examining navigation strategies as mice search for and approach an odor source. After mice were trained to navigate to odor sources paired with food reward, we observed behavioral changes consistent with localization 10-45cm away from the source. These behaviors included both orientation towards the source, decreased velocity, and increased exploration time. We also found that the amplitude of ‘casting,’ which we broadly define as lateral back and forth movement of the olfactory sensors, increased with proximity to the source. Based on these observations, we created a concentration-sensitive model to simulate mouse behavior. This model provided evidence for a binaral-sniffing strategy (inter-nostril comparison of concentration in a single sniff) and a serial-sniffing strategy (sampling concentration, moving in space, and then sampling again). Serial-sniffing may be accomplished at farther distances by moving the body and at closer distances by moving the head (casting). Together, these results elucidate key components of behavioral strategies for odor-based navigation.

**SUMMARY STATEMENT:** Use of a naturalistic odor-source localizing task uncovers key strategies underlying successful mouse navigation. Concentration-dependent models successfully recapitulate mouse behavior and reveal important behavioral components.

## INTRODUCTION

Mice, insects, and other animals navigate odor trails and locate odor sources with high fidelity, a remarkable feat given the complexity and variability of odor stimuli. Prior studies of olfactory navigation have focused on the mechanisms of navigating through odor plumes in air (Gire et al., 2016; Vickers, 2000) or water (Koehl et al., 2001), or of scent-tracking on the ground, specifically following odor trails (Khan et al., 2012; Means et al., 1992; Porter et al., 2007; Thesen et al., 1993; Wallace et al., 2002). However, different strategies may emerge for search under conditions near surfaces where intermittency (the fraction of samples above detection threshold) is high (Connor et al., 2018). In this study, we examine the odor-driven behaviors involved in the search for a discrete odor object under low wind conditions where gradient information is preserved, and develop a model of olfactory navigation to describe multiple strategies used for effective odor localization.

One general navigation strategy preserved across species and environmental conditions involves the use of lateral back-and-forth changes in orientation of the olfactory sensors during active odor-sampling (Baker et al., 2018). Moths navigating odor plumes use this strategy to reacquire an odor plume, a behavior termed “zig-zagging” or “casting” depending on the magnitude and angle of displacement (Carde & Willis, 2008; Kennedy, 1983; Kuenen & Carde, 1994; Marsh et al., 1978). Flies (Budick & Dickinson, 2006) and cockroaches (Willis et al., 2008) use a similar casting strategy during plume navigation. Lateral back-and-forth behaviors are also often observed in animals following odor trails. Rats employ head oscillations that sweep back and forth across a trail as they track it (Khan et al., 2012), a strategy also used by humans (Porter et al., 2007) and ants (Draft et al., 2018; Hangartner, 1969) following trails. In this study, we use the term ‘casting’ to describe the amplitude of the movement of the olfactory sensors during navigation toward an odor source (Baker et al., 2018; Zhao et al., 2017).

Successful navigation of a complex odor environment likely involves different behavioral strategies at different times. Knowledge of the odor environment, characteristics of specific odor cues, and general task context may each influence odor-guided behavior. Animals demonstrate strategy-switching between distinct behaviors as a consequence of both changes in odor stimulus properties and task context (Gire et al., 2016; Pierce-Shimomura et al., 1999; Thesen et al., 1993; Wesson et al., 2008). Environmental familiarity and visual cues also influence decision-making strategy during odor source localization. Given visual cues to navigate, exploratory casting based on odor cues disappears on tasks with a small number of potential goal locations; instead, mice begin directly navigating to known odor source locations in straight-line trajectories (Bhattacharyya & Bhalla, 2015; Gire et al., 2016). This resembles the transition from exploration to exploitation behavior observed during successive search trials (Hills et al., 2015; Sutton & Barto, 1998). Thus, both behavioral context and the availability of orienting stimuli are important for strategy choice in odor source localization.

Here, we use an open field odor-based spot-finding task to examine strategies of olfactory search for an odor source deposited on the surface under air flow conditions typical of an indoor setting. By randomizing odor source location, performing experiments in the dark, and using a discrete target odorant object, we prevent mice from switching to methods of systematic search based on memory, visual, or proprioceptive cues (Gire et al., 2016). We find that multiple dimensions of behavior are varied as mice approach the odor source, including velocity and casting, suggesting that these behaviors are associated with odor detection or localization. We then created agent-based models of olfactory search and use simulations to interrogate the roles of binaral-sniffing, variable velocity, and variable casting on effective odor localization. Both behavioral and simulation data suggest that multiple behavioral strategies are important for successful odor navigation to a deposited odor source under typical indoor air flow conditions in mice.

## MATERIALS AND METHODS

### Ethical approval

All experiments were completed in compliance with the guidelines established by the Institutional Animal Care and Use Committee of the University of Pittsburgh.

### Animals

Four male and one female adult M72-IRES-ChR2-YFP knock-in mice (The Jackson Laboratory) (N=5) were used for this study, with training begun after age P47. Average age at training start was P72. All mice were housed singly with behavioral enrichment (running wheel and house) and received unlimited access to water in their home cage.

### Behavioral arena

A large, open field behavioral arena was used for data collection, as described in Jones and Urban, 2018 (Figure 1A, adapted with permission from Jones and Urban, 2018). The arena is a custom built 36” × 45” transparent plexiglass and acrylic surface without walls mounted 4 feet above the floor in an aluminum frame. Rows of infrared light-emitting diodes (LEDs) were mounted along each of the four table surface edges. Data were collected using a camera mounted below the table surface (1280×1024 resolution, 11.2pixels/cm, 50 frames s^−1^, Flea3, Point Grey Imaging). Trials were run in the dark and LEDs from equipment in the room were blacked out with electrical tape. Remaining light from computer monitors was red-shifted so as to not be visible to the mice.

**Figure 1.**
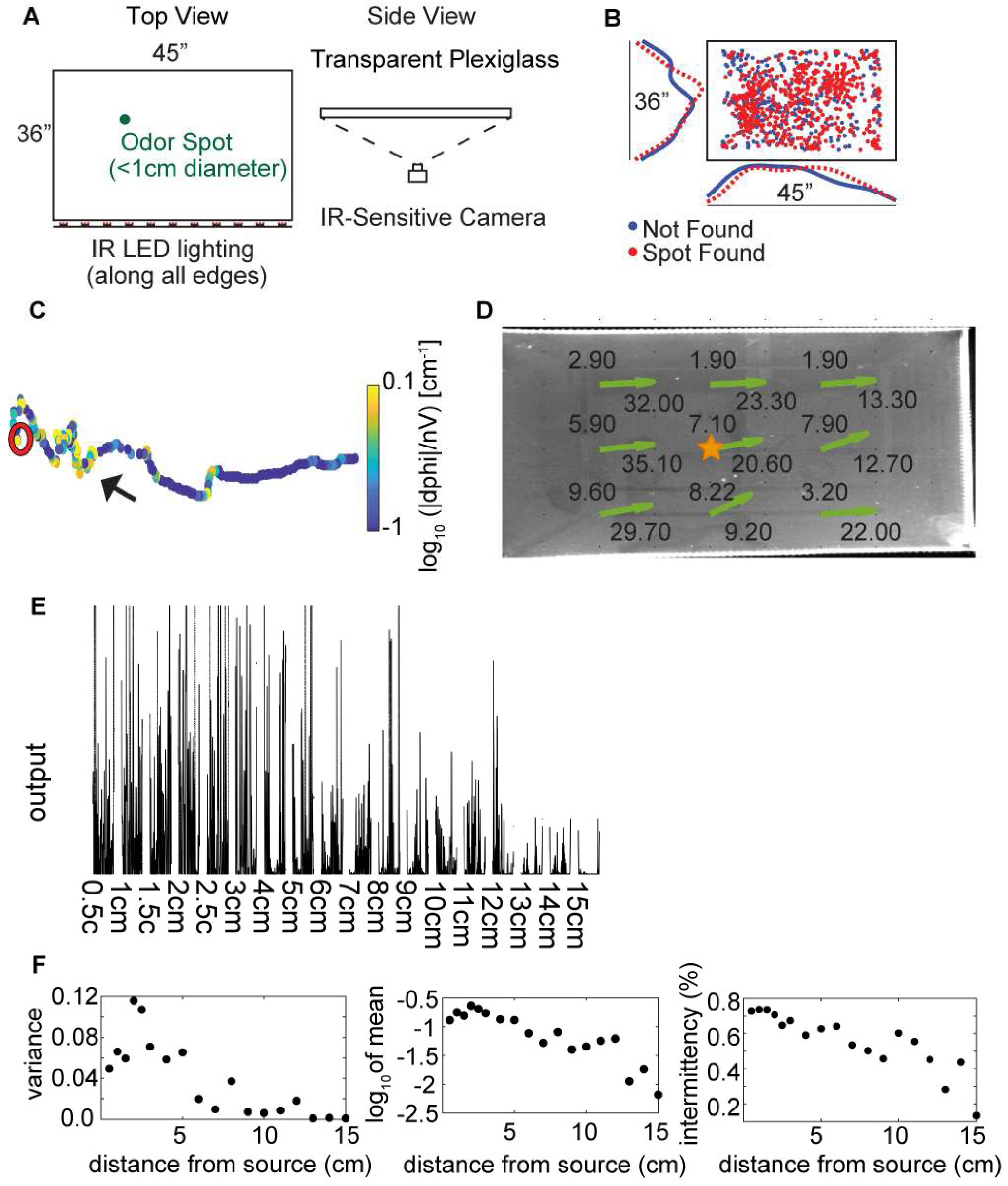
Experimental setup and methods. **A)** The behavioral arena is a 45” × 36” table made from three layers of plexiglass, with an infrared (IR) camera placed underneath to view the mice from below. Odor spots were placed pseudo randomly on the table by the experimenter before starting the trial, either unbaited or baited with a food reward. (Adapted with permission from Jones and Urban, 2018.) **B)** The pseudo random odor spot distribution for all trials (N=853). **C)** An example trajectory where the mouse finds the odor spot, showing nose tracking with the casting measure (unsigned curvature) on the color axis. The mouse is placed on the right side of the table and begins to hone in on the odor spot (red circle). The casting measure clearly indicates points of high curvature (yellow) from points of high linearity (blue). **D)** Anemometer measurements at 9 positions across the behavioral arena. Average wind speed in the <x> and <y> direction are indicated by the text at each point, with estimated wind velocity vectors indicated by the green arrows. **E)** Photoionization Detector (PID) readings from 0.5cm to 15cm away from an odor source placed at the center of the table (gold star in Figure 1D). Measurements were taken to the right of the star, in the ‘downwind’ direction. **F)** PID analysis suggests that mean signal variance, mean signal amplitude, and intermittency decrease starting at about 2.5cm from the source out to about 15cm.

### Odorized crayon

We created odorized wax crayons similar to what was described in Jones and Urban, 2018. Briefly, 4.9 g of a Crayola crayon, 0.1 g of Crayola chalk, and 0.1 g of odorant consisting of one of three different concentrations (0.1%, 1%, and 2% by weight) of methyl salicylate (Sigma-Aldrich) diluted in mineral oil (Fisher Chemical Paraffin Oil, 0121-1) were mixed together, and the mixture was poured into a silicon or rubber crayon mold. After ten minutes, the mixture cooled and hardened into an odorized crayon.

### Task

Prior to each trial, the table was wiped clean with 200 proof ethanol using a paper towel. After the table dried, a spot ~1cm in diameter was drawn in a pseudorandom location in the central area of the table (Figure 1B). Based on recent spot placements, the experimenter attempted to place the spot at a different location each subsequent trial. For baited trials, a small piece of chocolate or peanut (~0.1 g) was placed in the middle of the spot, while for unbaited trials, the spot was left empty. No more than 2 unbaited trials were run on a given day to prevent extinction of motivated behavior. With the lights off, a mouse was placed onto the table at a pseudorandom starting location along the right side of the table and allowed to freely explore the arena for approximately 3 minutes. Over the course of 5 months, 133-197 trials were collected per animal, 3-5 trials per day, 3-5 days per week, at a time between 9 am and 2 pm on a normal light-dark cycle.

### Food deprivation and task training

To motivate mice to perform the task, they were food deprived to 78±07% of original body weight. Animals received between 1.8-2.2 g of their normal chow per day and were given free access to water. Animals were trained to associate a spot of odor (<1cm diameter) with food reward, either a small sliver of peanut or milk chocolate. Training consisted of two stages over the course of 7-10 trials: 1) habituation (2-3 days); and, 2) odor-reward association (5-10 days). During habituation, mice were placed on the table for one 15 minute long trial on two consecutive days and allowed to explore the arena with no food reward present. During initial food-odor association, plexiglass dividers were used to decrease the arena size to 25% of the full arena. A baited odor spot was placed on the table. Once animals found the baited spot in the small arena during at least 66% of the day’s trials, the arena size was increased to 50% the size of the full arena, followed by an increase to the full 45” x 36” arena once the spot was again found during 66% of the day’s baited spot trials. Unbaited spot finding trials were introduced only after the animals successfully performed the baited spot task in the full arena. Training ended and data collection began upon successful performance of unbaited spot trials, which occurred following 6-13 days of training (18-39 trials). All reported data are from the full sized arena, but contain both baited and unbaited trials. Baited and unbaited trials were analyzed separately, but were found to be qualitatively similar and so are grouped together.

### Video Processing

44 trials were removed from analysis for technical reasons, leaving N=915 trials analyzed. Trials were removed if 1) the video file was corrupted; or, 2) tracking the mouse using Optimouse (see below) was judged by the experimenter to be unsuccessful before data analysis began. This likely occurred due to unintended variation in table illumination on different trials. Of these 915 trials, 54 had odor spot placement within 4 cm of the edge, and were removed from further analysis due to potential obstruction of mouse trajectories near the spots. Finally, 5 trials were removed because the mouse found the spot within 1 s, indicating that it was initially placed on the arena too close to the odor spot (most trials where this occurred the mouse started within 10 cm of the odor spot). This left N=853 trials analyzed here. Raw avi formatted files were processed using the Optimouse open source program (Ben-Shaul, 2017) to extract body position and nose position. Extracted position files were then post-processed with in-house software written in Matlab (www.mathworks.com). Position frames were then passed through a 5^th^ order median filter to suppress jitter in the position data and remove high-frequency noise. Position data was excluded if 1) Optimouse failed to detect a position, 2) the log_10_ of mean mouse brightness was less than 0.5 times the mean for that trial, 3) if mean velocity was greater than 100 cm s^−1^, 4) if the mouse length was greater than 8 cm, 5) if the mouse position ‘jumped’ by more than 35 pixels (3.13 cm) between consecutive frames, and 6) if the mouse position was 4 cm or closer from the edge of the arena. Note that the edges of the arena were metal fasteners and the mouse was often partially obscured from below when it walked around the edges. Exclusion criteria 6) accounted for the majority (N= 6E6; 84%) of excluded frames, as mice spent a good deal of time at the edges of the arena. Finally, a nearest neighbor filter was used to remove times when the mouse was stationary as the computation orientation of the mouse was unstable here. For the majority of analyses, we focused on either the time before a spot was found, or a matched sample of times for unsuccessful trials.

### Behavioral Analysis

A successful spot trial was scored when the mouse’s nose came within 1.5 cm of the spot, based on a sharp increase in the survival function on minimum distance to the spot (Figure 2A).

**Figure 2.**
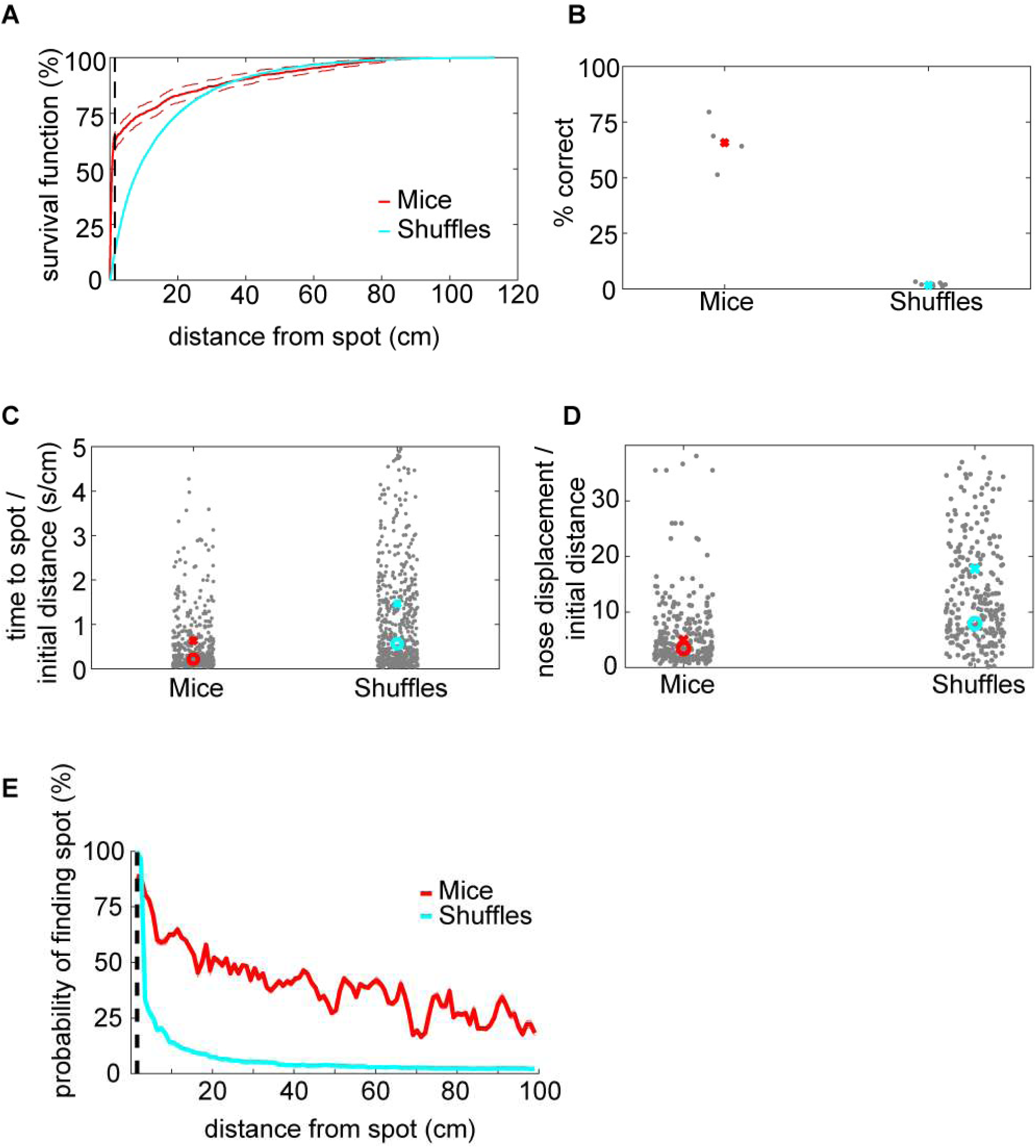
Mice and the CSM efficiently locate the odor source. **A)** The survival function shows the fraction of closest distances to the spot as a function of distance from the spot. At about 1.5cm away from the spot, the closest distance starts to drop off for mice, suggesting this would be a good threshold distance for counting a trial as successful. In comparison, there is no sharp distinction in the shuffled control data. Instead, shuffled minimum distances tend to increase smoothly as a function of increasing distance. At about 30cm away from the odor spot, the minimum distances are indistinguishable between mice and shuffled controls. **B)** Mice are more successful at finding the odor spots than shuffled controls, suggesting that they are using odor cues to guide their search. **C)** The time to spot is divided by the initial distance to the spot, and mice find the spot more quickly than shuffled controls. **D)** The nose displacement was divided by the initial distance to the spot, and the relative distance traveled by the nose of mice is less than that of the shuffled controls. **E)** The number of samples in 100 concentric rings around the spot were counted up. The number of samples in each ring on successful trials was divided by the total number of samples to give a measure of probability of finding the spot as a function of distance. This measure decreased roughly linearly for mice but highly nonlinearly for shuffled controls. This suggests mice are efficiently navigating toward the spot.

The fraction of the arena explored was calculated by binning the arena into approximately equal-area parts of 9-10,000 partitions and, then, the percentage of time exploring the center of the arena (excluding 4 cm around the edges) was calculated.

Analysis of percent exploration time entailed defining 100 circular rings around the odor spots at intervals of 1 cm. The number of samples in each ring was then normalized by the total number of samples in the trial to (% time). For trials when the mouse failed to find the odor source, the entire session was used.

To compute the probability of successfully finding an odor source as a function of distance, the occupancy was divided into successful trials (hits) and unsuccessful trials (misses). The fraction of hits was divided by the total number of trials (hits + misses) and averaged across mice.

The orientation relative to the odor spot was calculated by computing the difference in the four-quadrant arctangent of the vectors from body-to-nose and body-to-spot (Figure 3E).

**Figure 3.**
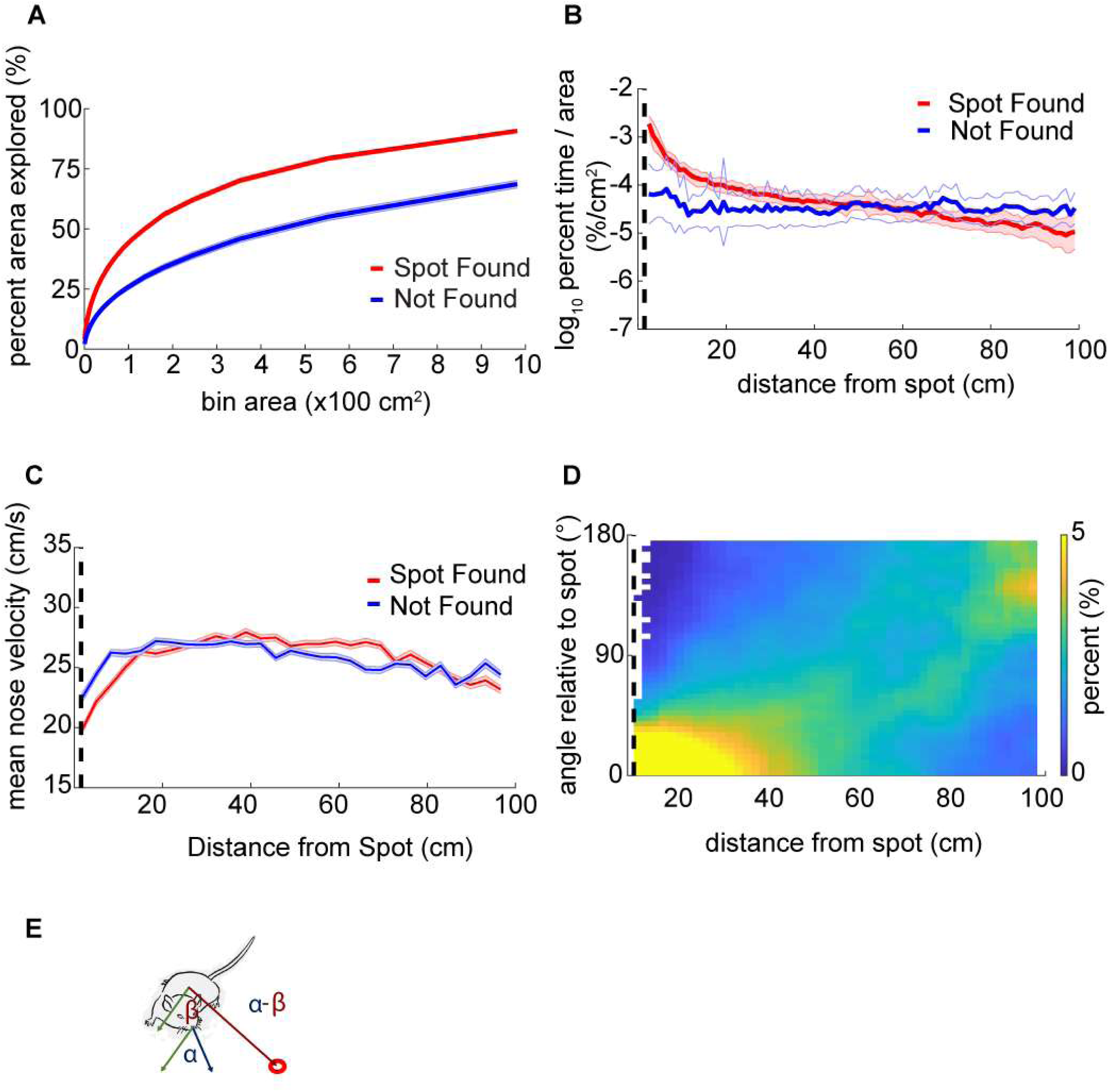
Mouse behavior varies systematically as mice approach the odor source. **A)** The center of the arena was broken up into bins of different area and the percentage of those bins visited by mice was counted up and plotted versus the bin area. In general, this ‘exploration percentage’ increased with increasing bin size, while successful trials entailed a higher exploration percentage than unsuccessful trials. **B)** The number of samples in 100 concentric rings around the spot was tallied, and normalized by both the total number of samples in a given trial and the area of the ring. This gave a measure of occupancy with units of %/cm^2^. As mice approach the spot on successful trials, their occupancy increases. In contrast, the occupancy on unsuccessful trials remains flat as a function of distance. **C)** Mean nose velocity decreases as mice approach the spot on successful trials. In contrast, mean nose velocity remains flat as a function of distance on unsuccessful trials. **D)** The orientation of the mouse relative to the odor spot was computed and normalized to 100% at each distance from the spot. For this measure, orientation of 0° indicates that the mouse’s nose is pointing directly at the odor spot and an orientation of 180° means that it’s nose is pointing away from the odor spot. Orientation tends to converge at 0° as mice approach the odor spot. **E)** Diagram demonstrating the computation of orientation relative to the spot. The angle β between the body and spot (red circle) is subtracted from the angle α between the nose and the body.

Derivatives were computed using the adaptive windowing Janabi-Sharifi algorithm (Janabi-Sharifi et al., 2000) available online (Manurung, 2016), but modified to handle missing data points (NaNs) and to allow for post-smoothing. Parameters used were a window size of 50 (1 s) and an error term of 0.2, with post smoothing of 0.1s.

Casting behavior was quantified by computing the numerical derivative of the arctangent of the nose velocity, taking the absolute value, and then dividing by the linear velocity of the nose (|dPhi|/nV). This is a modification of the zIdPhi measurement (Papale et al., 2012). |dPhi|/nV is equivalent to the signed curvature formula with units of m^−1^ and is therefore an appropriate measure of tortuosity. The algorithm was extensively tested on variable-frequency sine waves to verify that segments of high curvature were well-separated from segments of high linearity across a wide range of frequencies. Then, for position data from mice, the log_10_ of the absolute value of |dPhi|/nV was taken to generate a roughly normal distribution. An example of the |dPhi|/nV casting measure (Figure 1C) for one analyzed behavioral trial shows ‘elbows’ and angled segments with high curvature (black arrow) well-separated from straight segments of high linearity.

An alternate measure of casting used was the angle of the nose relative to the midline. For this measure, the angle from the nose to the midline was determined by calculating a vector for the mouse’s current heading and computing the shortest distance from the detected nose point to that line, then taking the arccosine of that angle.

### Anemometer Measurements

A hot wire anemometer (TSI Velocicalc 9535A) was placed at 9 different positions across the table (Figure 1D). 10 measurements were taken with the sensor aligned to the long-axis or short-axis of the table at each position and the respective averages were taken.

### PID Measurements

A cotton swab dipped in 99% Methyl Salicylate liquid was placed at the center of the table (Figure 1D, gold star) at approximately 0.5cm from the surface. Concentration was measured for 30s at distances from 0.5cm to 15cm using a Photoionization Detector (PID) (miniPid, Aurora Scientific). Raw measurements had noise peaks at 60Hz and 72Hz, broad spectrum high frequency noise, and substantial baseline drift. To remove the noise peaks, two 2^nd^ order Butterworth IIR notch filters were used. To remove high frequency noise, a (0.2s backward, 0.02s forward) moving mean filter was used. To remove baseline drift, a baseline estimation and denoising function was used (Duval, 2018). The denoised traces (Figure 1E) show large signal fluctuations and, generally, decreasing mean output with distance. Intermittency was calculated as the mean number of points greater than a threshold of 0.005.

### Shuffled Spot Control

As a control, spots were randomly shuffled and paired with tracking data from one session. Shuffled spots had to be at least 5cm away from the original spot. Analyses were repeated with the shuffled control data.

### Model Specification

Based on observations of mouse behavior, we developed an agent-based model that navigated a virtual odor environment. This agent made temporally discrete sniff-to-sniff comparisons of odor concentration as it moved through virtual space, altering its heading toward higher concentrations and away from lower concentrations. The agent consisted of a body with coordinates (*x, y*) and moved through space along a heading *θ* at a velocity *ν*_*c*_:

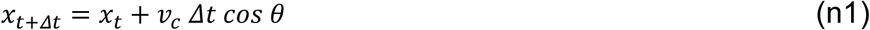

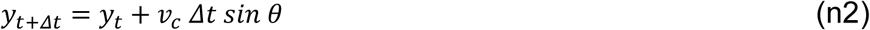

where Δ*t*=0.1 s represents a constant the inter-sniff interval. The agent’s nose was located at a distance *ℓ*=5 cm from the body at coordinates (*x*_*n*_, *y*_*n*_). The nose was further divided into left and right nares, (*x*_*nL*_, *y*_*nL*_) and (*x*_*nR*_, *y*_*nR*_), separated by an inter-nares distance, *d*_*nares*_=1.8 mm. The two nares are separated by an angle 2γ. The nose was capable of limited movement independent of the body heading, with an angular deflection *ϕ* relative to the forward heading. Nares positions were given as:

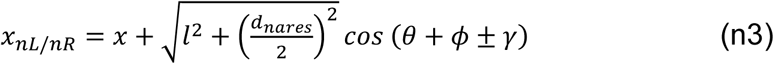

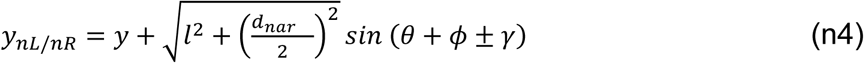

with nose deflection bounded within ± *ϕ*_max_ as dictated by mouse anatomy. The agent geometry is shown superimposed on a mouse body (Figure S2A).

The independent motion of the nose is what allows the agent to sample concentrations to the left and right of its current heading and adjust its heading toward higher concentrations (Figure 5C). The distribution of observed mouse nose deflections was approximately Gaussian (Figure S2C); therefore, we modeled the angular movement of the nose as an Ornstein-Uhlenbeck (OU) process, a mean-reverting correlated random walk with a Gaussian stationary distribution. At each sniff, the nose deflection was updated according to:

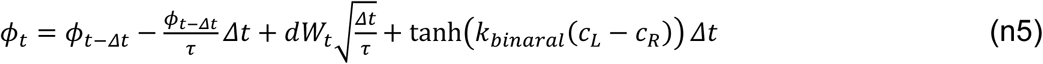

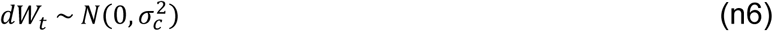

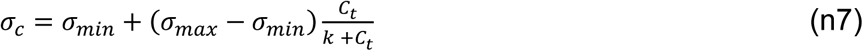

where *τ* is the characteristic time constant of the process, *σ*_*c*_ is the concentration-dependent standard deviation of the nose deflection, *dW*_*t*_ is a normally distributed random variable with mean of zero and standard deviation *σ*_*c*_. To this OU process, we add binaral bias in the form of the hyperbolic tangent of the left-right concentration difference; when *C*_*L*_ > *C*_*R*_, the nose is biased to the left and *vice versa*. Here, *k*_*binaral*_ controls the response of the nose to binaral inputs. The concentration value *C*_*t*_ is the average concentration across both nares at time *t*:

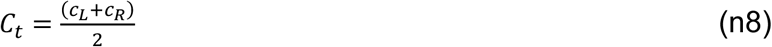

Here, *c*_*L*_ is the concentration at the left naris and *c*_*R*_ is the concentration at the right naris. We observed that mouse speed increases with distance from the odor source (see Figure 3F). Because the agent in our model does not have information about its distance from the odor source, instead we used a concentration-dependent modulation of speed. Specifically, we use a sigmoidal function with a quartic coefficient to approximate concentration-dependence of speed:

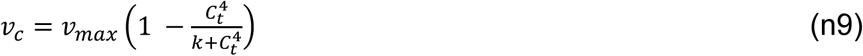

where *v*_*max*_ is the maximum velocity and *k*=½ concentration units. Note that this is the same *k* that appears in eq n7.

**Figure 5.**
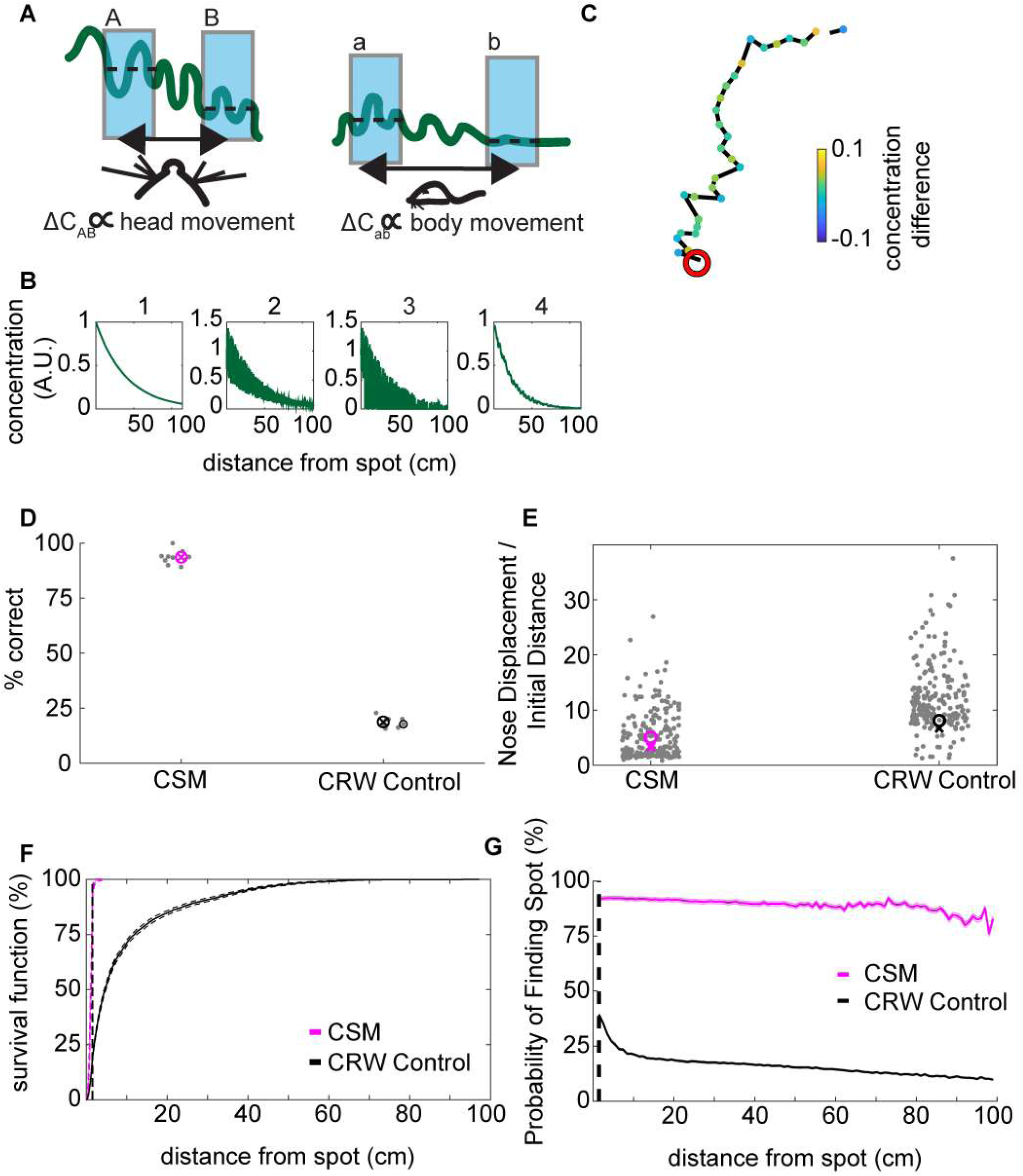
Development of concentration-sensitive model. **A)** Our general hypothesis in constructing the model was that lower concentration differences (ΔC_ab_) far from the source would be detectable by gross movement of the body (right panel). In contrast, higher concentration differences (ΔC_AB_) would be detectable by fine movement of the head (left panel). In this illustration, the wavy green line represents the concentration of an odorant. The blue shading indicates averaging concentration over an inhalation. The dashed lines indicate the mean concentration for each sniff. **B)** The odor concentration profile was constructed in four steps: 1) A smooth concentration gradient was computed; 2) multiplicative random noise was added to the gradient; 3) ‘Intermittency’ was modeled by setting a proportion of pixels equal to zero. This proportion increased with distance from the source. Finally, 4) Gaussian smoothing was implemented to simulate averaging of the signal over a sniff. **C)** An example trajectory from the concentration-sensitive model (CSM) that successfully found the odor spot (red circle). Concentration difference (A.U.) is indicated. **D)** The success rate of the CSM was above 75%, while the CRW control model had a success rate of below 25%. **E)** The nose displacement divided by the initial distance to the spot was lower for the CSM than for the CRW control, indicating more direct travel to the odor spot using concentration to guide the model. **F)** The survival function of approach distances shows that nearly all CSM trajectories approach within 30cm of the source, while CRW control trajectories are more spread out across the entire table. **G)** The probability of finding the odor spot was computed for the models as in Figure 2E. With this measure, the CSM was more successful at finding odor spots at all distances from the spot compared to CRW controls.

### Model Odor Environment

The agent must iteratively sample concentrations in an odor environment to navigate in the direction of higher concentration. To simulate the ambient concentration, *C(x,y)*, we use a noisy bivariate exponential distribution normalized to a value of one at the source:

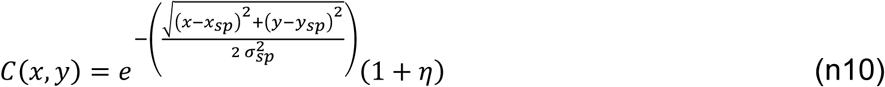

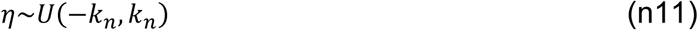

Here, (*x*_*sp*_, *y*_*sp*_) is the location of the source, *σ*_*sp*_ controls the width of the odor distribution around the source (model results were obtained with *σ*_*sp*_=20 cm), and *η* is multiplicative uniform noise added to each point in a spatial grid to approximate the effect of turbulent flow, bounded at *k*_*n*_=0.5.

To represent the table, we generate a static smooth odor landscape grid with a resolution of 1 mm according to eq n10 (Figure 5B, step 1). Next, we then apply noise according to eq n11 (Figure 5B, step 2). We then simulate the PID intermittency in the odor signal by randomly setting grid values to zero (Figure 5B step,3). The probability that a grid value is zeroed, *P*_*int*_, increases with distance from the source according to

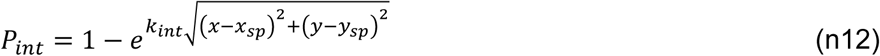

where *k*_*int*_ is the decay parameter set to −0.002 cm^−1^. Finally, we apply a Gaussian filter with a standard deviation of 4 mm to introduce local spatial correlations into our landscape (Figure 5B, step 4).

To reflect the apparent lack of mouse orientation toward the source beyond 30 cm, (Figure 3D), the models have a concentration detection threshold of 0.25 concentration units. This represents a distance of approximately 30cm from the source.

A distribution of simulated spot positions was used for simulations, matched to data (Figure 1B), but due to edge effects reducing overall model performance, spots closer than 10cm to the walls were not used.

### Concentration-Sensitive Model (CSM)

The agent moves along forward heading *θ* at velocity *v*_*c*_, iteratively sniffing the concentration at its nose coordinates (Eqs n1-n4). At each sniff, the agent’s nose randomly samples to the left or right of the forward heading depending on the state of the random nose deflection process (Eqs n5-n7). It compares the current concentration sample, *C*_*t*_, to the sample from the previous sniff, *C*_*t-Δt*_. If *C*_*t*_ is greater than *C*_*t-Δt*_, the agent sets its heading to the direction of the nose, *θ*_*t+Δt*_ *:= θ*_*t*_ *+ ϕ*_*t*_. If *C*_*t*_ is less than or equal to *C*_*t-Δt*_, the agent turns away from the direction of the nose, *θ*_*t+Δt*_ *:= θ*_*t*_ *-ϕ*_*t*_. This algorithm is effective at reproducing key parameters of mouse odor source localization (Figure 5):

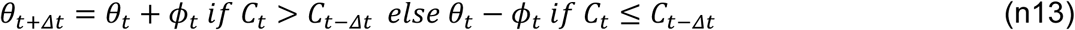

### Correlated Random Walk (CRW) Controls

To test model behavior in the absence of odor-sensitivity, and to serve as a control in case mice were randomly finding the spots, concentration-dependent turning was removed from the model and replaced by a random turning decision. Eq n13 is replaced by:

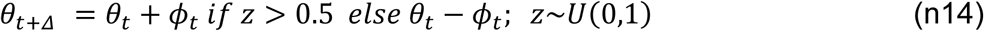

To our surprise, CRW control models had a similar success rate to mice when run for 180 s. After observing mice on videos; however, we determined they were not exploring the arena for the entire 180 s trial. On successful trials, before finding the odor spots, the average time mice actually spent exploring the center of the arena was only ~30 s. The CRW controls continuously explored the center of the arena as their decision rule at table boundaries was to bounce off like a billiard ball. Therefore, we ran all simulations for 30s to match average mouse exploration time in the center of the arena.

### Reduced Model Controls

To evaluate the effects of different hypothesized features of mouse navigation, we constructed 8 model variant combinations by leaving out binaral-sniffing, concentration-dependent velocity, and/or concentration-dependent casting amplitude. We then evaluated CSM model performance in the absence of these features. To eliminate binaral-sniffing from the model, *k*_*binaral*_ in eq n5 was set to zero. To remove concentration-dependent velocity from the model, eq n9 was modified so that *v*_*c*_ was equal to the constant *v*_*max*._ To remove concentration-dependent casting amplitude from the model, eq n7 was modified so that *σ*_*c*_ was set equal to the constant *σ*_*min*_. The same 8 model variants were created for the CRW control, and results were averaged together to obtain a control distribution (Figure 6, black lines; Figure 7A, grey bar).

**Figure 6.**
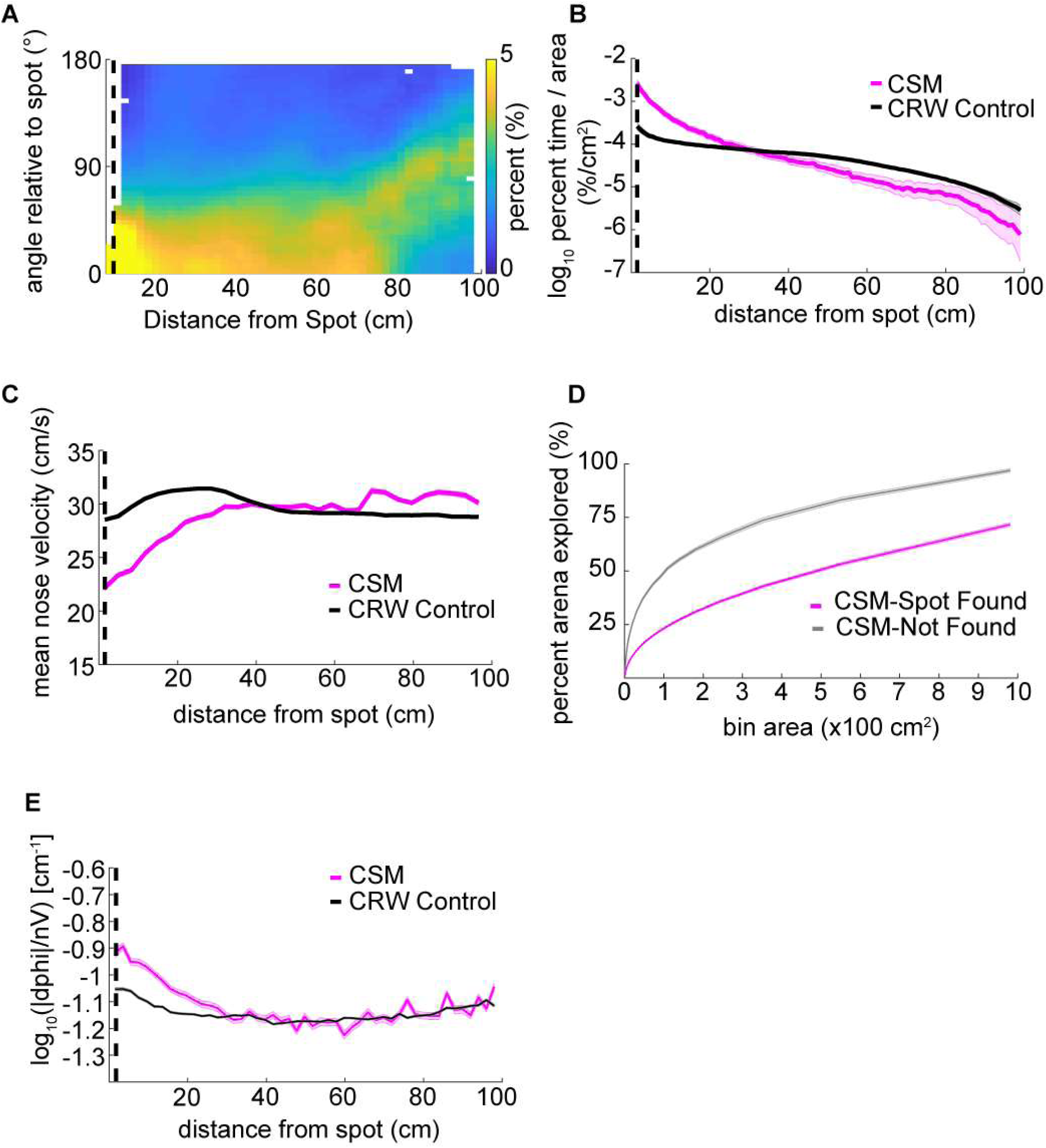
Detailed analysis of CSM trajectories. **A)** The orientation of the CSM model relative to the odor spot resembled that of mice (compare Figure 3D). **B)** The occupancy of the CSM resembled that of mice (magenta line; compare red line Figure 3B), and significantly diverged from the CRW control (gray line) at around 20cm from the odor spot. **C)** The mean nose velocity of the CSM (magenta line) decreased as the model approached the odor spot, consistent with what was observed with mice (compare red line Figure 3C). The nose velocity of the CSM was different from the CRW control (gray line). **D)** In contrast to mice, successful trials of the CSM (magenta line) showed *less* exploration than unsuccessful trials (gray line) (compare red line to blue line in Figure 3A). **E)** Casting increased as the CSM approached the spot (magenta line) consistent with observations in mice (compare red line Figure 4A). Casting in the CSM increased with respect to the CRW control (gray line) at a distance of 7cm.

**Figure 7.**
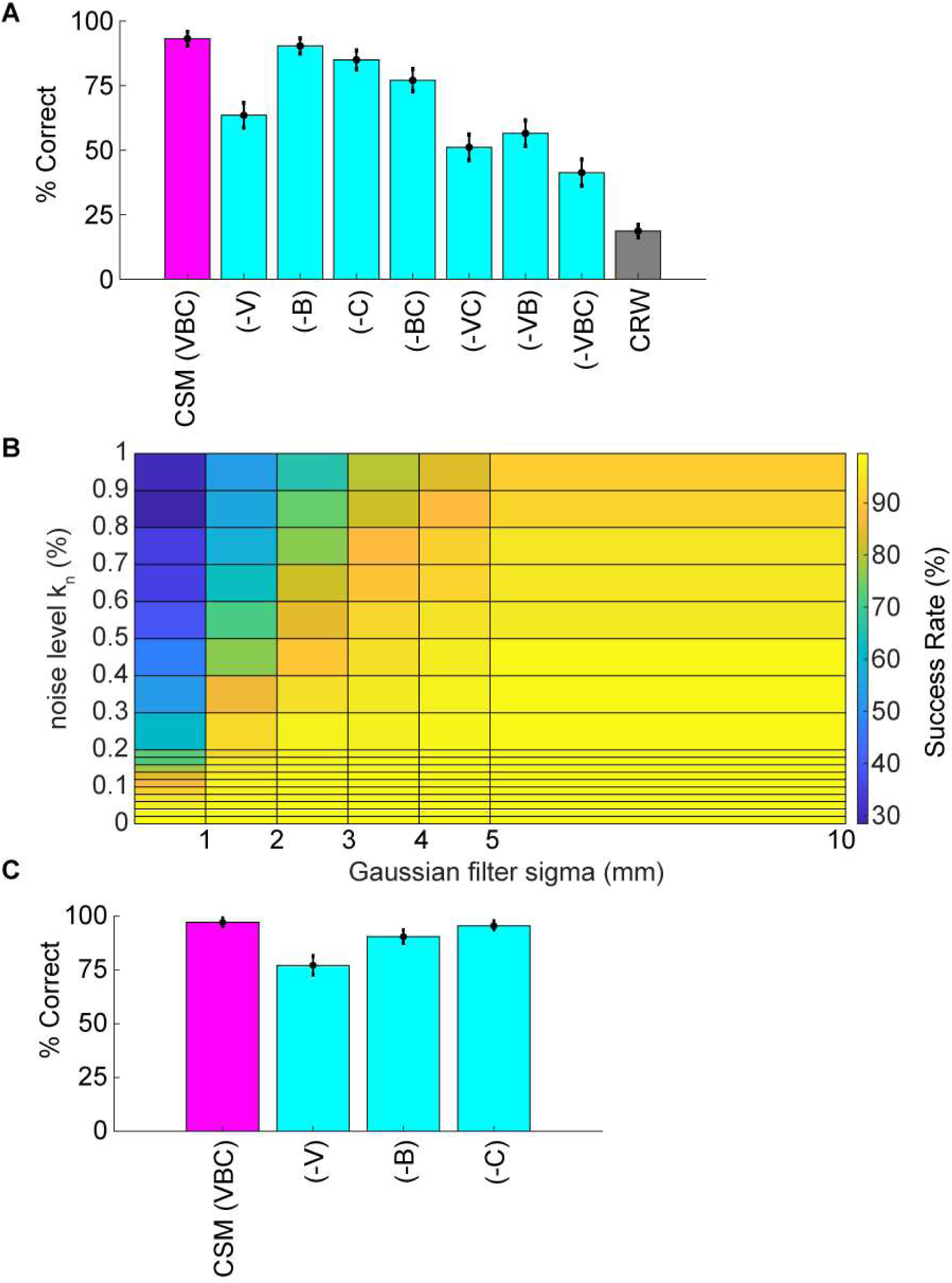
Analysis of reduced models and parameter sweeps. **A)** The full model (CSM) included three components: 1) a concentration-dependent velocity term (V), 2) a concentration-dependent casting term (C); 3) a binaral-olfaction term (S). Removal of any of these components (-V), (-B), or (-C) may, in principle, reduce model performance. Removal of concentration-dependent velocity (-V) had the largest effect on performance. Removal of concentration-dependent casting (-C) had a smaller effect on performance. Removal of binaral-olfaction (-B) had the smallest effect on performance. Removal of both concentration-dependent velocity and casting (-VC) had a multiplicative effect on performance. Removal of all three components (-VBC) was still significantly better than CRW controls. **B)** Comparison of noise effects on CSM success rate shows that the CSM model is robust for a wide array of different noise conditions. **C)** Because (-B) had the smallest effect on performance, we wished to investigate its role further. We added a coefficient in front of the hyperbolic tangent term in eq n5 and set it equal to 3. With this additional term, success rate reduction from removing the binaral term increased. However, success reduction from removing (-V) or (-C) terms decreased. This suggests that binaral sniffing can compensate somewhat for lack of velocity or casting changes.

## RESULTS

### An open field odor-based spot-finding task was developed

To examine the behaviors relevant to odor source localization, we designed a task that minimized non-olfactory cues. Five mice were trained to find a <1cm diameter spot of odorized wax within a rectangular behavioral arena of approximately 1m^2^ in the dark (Figure 1A, and Methods). Odor sources (methyl salicylate with(out) bait) for each of the trials (N=855) across all mice (N=5) were pseudo randomly distributed within the center of the testing table (Figure 1B). An example trajectory shows the nose position of the mouse starting at the right-hand side of the arena and moving leftward before successfully finding the spot (red circle; Figure 1C). In this figure, the color indicates a measurement of curvature (|dPhi|/nV; see Methods) used to detect casting amplitude. Additional example trajectories are displayed in Figure S1, with color indicating relative time (in seconds).

### Wind Conditions in the Room

We did not generate any airflow during these experiments but of course air flow occurs in every typical environment and air movement likely dominates the effects of odor diffusion in the vast majority of environments. The room in which these behavioral data were acquired had a ventilation input vent at roughly the center of the room and the primary outflow in the room was underneath the door. Airflow measurements were taken on the behavior table (height 1.135m above the floor) at 9 positions and 2 orthogonal orientations. Wind speed was measured between 1.9ft/min – 9.6ft/min along the short axis and 9.2ft/min – 35.1ft/min along the long axis of the table (Figure 1D). Wind direction was estimated by taking the four-quadrant arctangent (green arrows).

### Photoionization Detector (PID) Measurements on Behavior Table

A cotton swab dipped in 99% methyl salicylate was placed at the center of the table (Figure 1D, gold star). The tip of a PID was placed at distances from 0.5cm to 15cm away from the odor source. After denoising (see Methods) PID traces were highly variable but mean output decreased reliably with distance (Figure 1E). Three aspects of the PID data were characterized. The mean variance increased for distances less than 2.5cm and then decreased (Figure 1F, left panel). The increase at small distances could be due to saturation of the PID at high concentrations. The mean output also decreased with distance from the source from about 2.5cm (Figure 1F, center panel). Intermittency was measured as the fraction of samples above baseline, and decreased with distance from the source (Figure 1F, right panel).

### Mice learn the open field odor-based spot-finding task

To determine an appropriate distance threshold for achieving a successful trial and an approximate minimum concentration-detection threshold, we examined the survival function with respect to distance from the spot (Figure 2A). Typically, a survival function is used to determine the probability that a variable of interest survives past some time point; here, we use it to describe the probability of a mouse reaching a given distance away from the odor spot. For mice, there was a sharp increase in survival at about ~1.5 cm, with a more gradual increase toward 100% at about 90 cm from the odor spot (red line). Thus, mice on all trials successfully came within ~90 cm of the spot. In contrast, the survival function from shuffled data rose smoothly, crossing the mouse data at about 30cm. We therefore set the distance threshold for finding a spot to be 1.5cm and estimated the minimum concentration-detection threshold to be 30cm for later modeling purposes.

Given a threshold of 1.5cm, mice were successful in 64% (N=532/826) of trials (Figure 2B). In comparison, shuffled controls were successful a mere 1.67% (N=1,524/91,446). Mice had a significantly higher success rate than shuffled controls (Welch’s t-test; p=6.22E-5, t-stat=14.69, df=4). A logistic regression on successful versus unsuccessful trials for mouse data revealed several highly predictive explanatory variables (Table S1). Baited trials had higher success rates (p=4.68E-12, β=2.51), potentially indicating higher motivation during these trials or a contribution of the odor of food to the localization. Greater arena exploration also predicted success (~45 cm^2^ bins; p=1.65E-15, β=15.51; see Figure 3A). Slower mice (p=6.89E-06, β=-0.29) were more successful, though they had faster average nose velocity (p=1.07E-04, β=0.27) and higher casting (integrated log_10_[|dphi|/nV] within 30cm; p=2.56E-3, β=1.4), suggesting that slower forward body velocity and higher casting amplitude may be features of successful search strategies. Successful completion of this spot-finding task is thus dependent on features of individual trajectories.

We conclude that mice find odor spots above chance levels established by the shuffled control, and predict that further evaluation of specific trajectory features will provide useful information about odor localization strategy.

### Mice efficiently navigate to the odor source in comparison to shuffled controls

Next, we compared navigation efficiencies of mice to data from shuffled controls by examining time to spot divided by initial distance, nose displacement divided by initial distance, and the probability of finding the spot as a function of distance. Time to spot was divided by initial distance to spot, with the average for mice (0.51 s/cm) significantly lower than shuffled controls (1.71 s/cm; Figure 2C; Welch’s t-test; p=7.25E-18, t-stat=8.72, df=1700). Then, nose displacement was divided by initial distance to spot, with the average for mice (4.75x) significantly lower than shuffled controls (17.78x; Figure 2D; Welch’s t-test; p=1.2E-19, t-stat=-9.11, df=1573). To determine the probability of finding the odor source at various distances away, successful samples (spot found) were divided by total samples (spot found + not found) in concentric annuli centered at the spot (Figure 2E). For mice, this resulted in a fairly linear distribution (red line), while for shuffled controls the distribution was highly nonlinear (cyan line). We conclude that mice are more efficient at navigating to the odor source than shuffled controls.

### Mouse behavior varies systematically as a function of distance from the odor source

As suggested by the logistic regression analysis of mouse data, exploration percentage, velocity, and casting amplitude are important for odor source localization. Next we tested whether mice reliably change behavior depending on proximity to the odor source. We conducted four analyses to test this hypothesis: 1) looking at the percentage of the arena explored, 2) computing the percentage of time exploring at a given distance on a given trial; and separately, 3) examining mouse velocity as a function of distance from the odor source; and 4) analyzing mouse heading relative to the odor source.

Using a bin size of ~45 cm^2^, mice explored an average of 33±17% (mean±s.d.) of the table before finding the spot. This estimate varied as a function of bin size and successful spot-finding (Figure 3A; two-way ANOVA interaction term; p<1E-100, F=83.95, df=97). Mice also explored progressively less of the table with increasing trial (~45 cm^2^ bins; ANOVA; p=2.65E-51, F=57.16, df=5), suggesting refinement of learning throughout the experiment.

We then computed the percentage of time mice occupied different distances from the odor source, normalized by the area of the annulus (Figure 3B). This revealed a decreasing function with distance on successful trials (red line) compared to a flat distribution with distance on unsuccessful trials (blue line). These lines diverged at a distance of 44cm (two sample Kolmogorov-Smirnov test; p<1E-10), suggesting that mice are exploring more within 44cm of the odor source on successful trials.

Next, we examined mouse velocity as a function of distance from the odor source (Figure 3C). As mice approached the odor source across all trials, nose velocity slowed down (ANOVA; p=2.72E-91, F=18.73, df=28), as did body velocity (ANOVA; p=1.02E-160, F=31.12, df=28).

We next examined mouse heading relative to the odor source for successful trials (Figure 3D). If a mouse detects odor cues that provide insight into the location of the odor source, then the mouse’s nose should orient towards the source. At each position sample, the difference in angle between body-to-spot and body-to-nose was computed (Figure 3E). We found that orientation relative to the spot depended on distance from the odor spot (Watson-Williams Test; p<1E-100, F=877.94, df=48), and has a median value not significantly different from zero (i.e. oriented directly at the spot) within 9.54cm (p>1E-3, circular test for equal medians with Bonferroni correction).

In summary, we found that mice vary their search strategy and orient towards the spot as they approach it. Mice also slowed down as they approach the spot. Together, these behavioral changes potentially represent a speed-accuracy trade-off during the final stage of navigation towards an odor object, roughly within ~ 10-45cm.

### Casting increases near the odor spot and is modulated by velocity

Casting is seen in diverse odor navigation circumstances ranging from plume navigation by moths to trail tracking by dogs; thus, mice may also cast during the open field odor-based spot-finding task. To measure casting amplitude, we calculated the log_10_ of the absolute value of the angular velocity of the nose divided by the linear velocity of the nose (Figure 1C; and see Methods). This is an appropriate measure for unsigned path curvature with units of (m^−1^). We found that curvature increased as mice approached the odor source (ANOVA; p=2.2E-117, F=23.4, df=28) with an interaction between distance and successful versus unsuccessful trials (two-way ANOVA; p=0.0041, F=1.85, df=28).

We used a stepwise Generalized Linear Model (GLM) to determine the relative effect of velocity and distance on the curvature measure of casting. The first term to include in the stepwise GLM was velocity (p<1E-100, F=9636, Deviance=1500) and the second was distance (p=7.1E-41, F=1481, Deviance=1481). No nonlinear terms were added to the model.

As curvature explicitly depended on inverse nose velocity (see Methods and results from stepwise GLM), we computed the angle between the head and body centerline as an alternate measure of casting (Figure S3B). Using this alternate measure of nose deflection, we again found that casting increases as mice approach the odor source (ANOVA; p=2.18E-56, F=12.82, df=28), with an interaction between distance and successful versus unsuccessful trials (two-way ANOVA; p=0.02, F=1.61, df=28).

To assess whether animals began using non-odor cues to navigate towards the odor spots on later trials, we performed one-way ANOVAs on curvature versus binned trial number (N=6). We found an effect of session on curvature (ANOVA; p=1.42E-11, F=12.34, df=5), suggesting that after 60-70 trials, a shift in strategy occurred which led to systematically less casting. Further analysis revealed that this effect was consistent with systematically increased nose velocity on later trials (ANOVA; p=9.16E-7, F=7.36, df=5).

We also examined casting amplitude as a function of approach vector. To compute approach vectors, the angle of the nose with respect to the body was taken and positions were centered with respect to the spot, then binned every 2.3cm. The length of the vector was then normalized by the standard deviation of the angles at that point. The vector colormap indicates the log_10_(|dphi|/nV) at that point. Note that on most trials mice approached the spot from the northeast direction (the ‘downwind’ direction, see Figure 1D). It was in this direction, at points closest to the source, where casting increased (Figure 4B).

**Figure 4.**
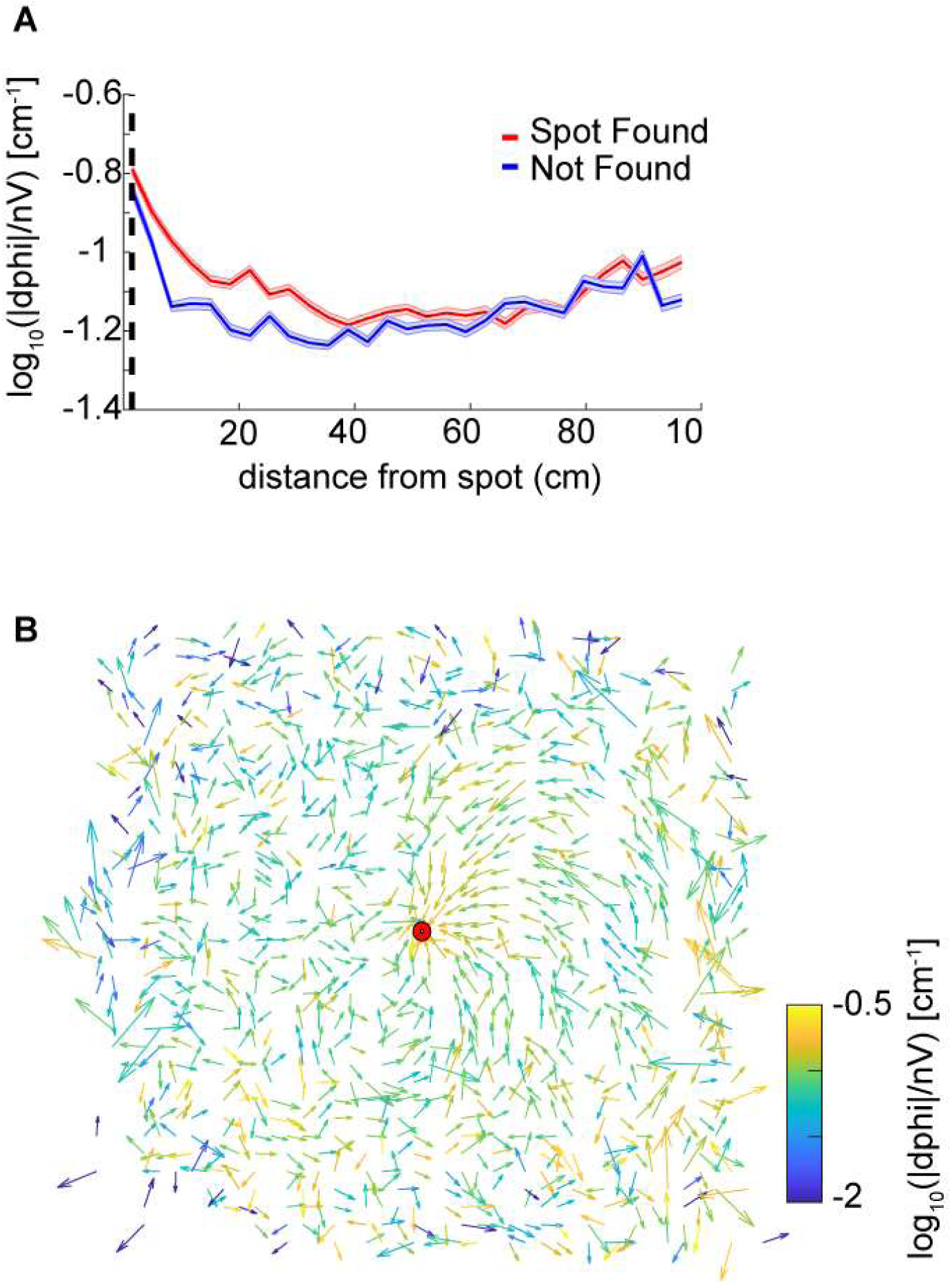
Analysis of casting amplitude using curvature. **A)** Unsigned curvature of the nose was computed as a measure of casting amplitude (see Methods) and averaged across successful trials (red line=mean±s.e.m.). Casting increased as mice approached the odor spot (ANOVA; p=1.21E-64, F=42, df=8), and was significantly higher on successful trials within ~12 cm compared to unsuccessful trials (Welch’s t-test, p=7.2E-5, t-stat=-4.26, df=61, Bonferroni correction). **B)** The approach vector for successful trials are the vector components of the circular mean of the nose-to-body orientation after centering position data at the odor spots. Vectors are normalized by the circular standard deviation. Color indicates the curvature measure for casting at that point. Most trajectories approach the spot from the northeast.

Overall, we found that casting amplitude of mice increases at distances close to the spot, suggesting that it may play a role in the successful navigation to a discrete odor object by mice. This change in casting is potentially correlated with velocity, which also decreases significantly during the approach to the spot.

### A concentration-sensitive model (CSM) was developed to capture key aspects of mouse behavior discovered on the spot-finding task

The core of the CSM model is the temporal comparison of sniff-to-sniff concentration. At each discrete “sniff,” it samples the odor at the position of its nose and compares the current odor concentration with the previously sampled concentration. Part of the algorithm averages across the two nares (Eq n8) while one component (binaral component) compares the difference between nares (right-most term in Eq n5). A typical simulated nose trajectory showing a successful capture displays inter-sniff differences in odor concentration in arbitrary units (Figure 5C). The CSM model used here had parameters: *Δt*=0.1s, *ℓ*=5 cm, *ν*_*max*_=25*cm/s*, k=0.5 concentration units, *τ*=0.1 s, *σ*_*min*_=0.2 radians, *σ*_*max*_=0.3 radians, and *k*_*binaral*_=200 The value of *τ* chosen gave a good balance of sharp turning nose trajectories with smooth nose trajectories. Values an order of magnitude lower resulted in tight spirals of the nose, while values an order of magnitude higher resulted in loss of maneuverability of the model. *ν*_*max*_ was chosen as it was the approximate average velocity of mice. Because simulated odor concentrations were scaled to a maximum of one, *k*=0.5 was chosen as the half-maximal value for behavioral response. Changing *k* could be used to ‘tune’ the model’s performance without qualitatively affecting its trajectories. Values of *σ*_*min*_, *σ*_*max*,_ and *k*_*binaral*_ were chosen so that the model’s nose curvature remained qualitatively “mouselike” in character. *I.e.*, the model trajectories did not appear overly tortuous or linear when compared by eye to the mouse trajectories.

To model the observed decrease in velocity close to the odor source, we implemented a concentration-dependent velocity term (Eq n9). In this term, the maximum velocity, *ν*_*max*_ decreases with increasing concentration according to a sigmoidal function. This function was chosen as it was roughly linear in the center of the concentration range and saturated at high concentrations. To model the increase in curvature with approach to the spot, a concentration-dependent widening of the normal distribution was implemented (Eq n7). These equations were linked by a common parameter *k* that controlled their rates of change as a function of concentration.

As a control, we created a Correlated Random Walk (CRW) model that turned independently of concentration (Eq n14), aligning its heading with its nose 50% of the time and away from its nose 50% of the time, resulting in left and right turns in equal proportion. Models using odor cues to navigate should outperform the CRW.

### CSM simulations have similar general properties to mice

From our data, we formulated a hypothesis that mice are detecting concentration gradients in order to navigate to the target. While some prior work has suggested that animals can navigate to sources without concentration gradients (Balkovsky & Shraiman, 2002; Vergassola et al., 2007), which may be necessary in highly turbulent environments, odor concentration gradients will develop over time through convection-diffusion and are a likely candidate for rapid olfactory navigation to targets under conditions of low turbulence. We refined our hypothesis to suggest that mice are 1) moving rapidly where the slope of the gradient is low and detecting changes at different physical locations (a versus b); versus, 2) moving slowly where the slope of the gradient is high and detecting changes by sweeping their heads back and forth (A versus B), which we interpret as casting (Figure 5A).

Initially, we had used a simple stimulus model of multiplicative random noise added to a smooth gradient. However, after examining the fluctuations of concentration near the table on a PID, we decided to implement a more complex odor stimulus model. This included 1) generation of a smooth concentration gradient; 2) addition of multiplicative noise which increased near the source; 3) setting pixels equal to zero at a proportion dictated by an intermittency fit such that the farther away from the source the higher proportion of zeros; and, 4) the smoothing of the resulting stimulus by a two-dimensional local Gaussian filter (Figure 5B).

The CSM model with 4-step stimulus generation generally resembled mouse behavior (Figure 5C). Here, concentration difference between consecutive samples is plotted so that the turning away from high-to-low and turning towards low-to-high concentrations is visible. The simulated odor spot is indicated by the red circle.

Spots positions for the CSM and CRW were taken from the mouse data, with spots less than 10cm from the edges being removed. The success rate of the CSM was 94%, significantly greater than the 19% for CRW controls (Figure 5D; Welch’s t-test; p=2.16E-22, t-stat=59.45, df=17). The CSM therefore successfully used information about concentration to navigate within 1.5cm of the odor source.

The nose displacement over initial distance ratio for the CSM (~5.6x) was significantly less than the CRW (~13x) (Figure 5E; Welch’s t-test; p=3.5E-11, t-stat=-34-41, df=9), indicating that the CSM found the odor spots more effectively than the CRW. The survival function of CSM minimum distance to the spot was shifted to left and up compared to the CRW control (Figure 5F). The probability of finding the spot measure for the CSM was also consistently higher than the CRW control (Figure 5G).

### Examining the structure of CSM trajectories suggests that they approximate mouse behavior

The orientation relative to the spot for the CSM was qualitatively similar to mice (Figure 6A, compare with Figure 3D). The CSM occupancy increased with respect to the CRW occupancy as agents approached the spot (Figure 6B; two-sample Kolmogorov-Smirnov test; p<1E-10 at <26cm). This was similar to the occupancy observed in mice on successful trials (Figure 3B). Mean nose velocity of the CSM decreased as agents approached the spot (Figure 6C), as expected, given eq n9. On successful trials, the CSM explored *less* than it did on unsuccessful trials (Figure 6D). This contrasts with the behavior observed in mice (Figure 3A), suggesting that there are unexplored differences between the CSM and mice. CSM casting increased as agents approach the spot (Figure 6E), as expected, given eq n7. This approximates the effect observed in mice (Figure 4A).

### Simulations allow separation of component strategies for odor-driven navigation

Simulations allowed for further dissection of the odor-driven factors that impact behavior and successful odor source localization such as decrease in velocity close to the source, increase in casting close to the source, and binaral sniffing. We created reduced CSMs to examine the relative contributions of three hypothesized behavioral components: binaral-sniffing, concentration-dependent velocity, and concentration-dependent casting amplitude. These model variants are denoted (-V) for the model lacking concentration-dependent velocity, (-B) for the model lacking binaral olfaction, (-C) for the model lacking concentration-dependent casting, (-VC), (-VB), (-SC) for models lacking multiple components, and (-VBC) for the model lacking all three components. While all reduced model variants outperformed CRW controls (Figure 7A), removal of any of these behavioral components decreased model success. The removal of concentration-dependent velocity led to the largest change in success rate (-V, Δ30%), followed by concentration-dependent casting amplitude (-C, Δ8%), and binaral-sniffing (-B, Δ3%). The change in success rate was significant for removal of concentration-dependent velocity (Welch’s t-test; p<1E-100, t-stat=169, df=1472), concentration-dependent casting (Welch’s t-test; p<1E-100, t-stat=58.3, df=1759), and binaral sniffing (Welch’s t-test, p=2.65E-100, t-stat=22.6, df=1927). The effect size depended on the number of bootstrapped samples (out of 100,000) and groups in each sample. For this analysis, we chose 1000 groups with 10,000 samples in each group, selected without replacement.

These behavioral components of olfactory search are synergistic; simultaneous removal of two or more of them leads to a greater reduction in success rate than removal of individual components. Model variant (-VC), in which both velocity decrease and casting increase are removed, resulted in a significant Δ12% change in model success relative to (-V) alone (Welch’s t-test; p<1E-100, t-stat=-55.9, df=1997). All groups performed significantly better than the CRW Control, even with all three additional components of the model removed (-VBC; Welch’s t-test; p<1E-100, t-stat=67.08, df=1997). This is logical since (-VBC) still makes decisions based on inter-sniff concentration differences, while the CRW Control makes random decisions.

Significance of model parameters was also established by shuffling 12 parameters of the model 50k times, and comparing the effect of each parameter using a logistic regression (Table S2). We also included two dependent variables from the model trajectories that we thought might impact success rate, the mean casting (log_10_(|dPhi|/nV) within 30cm, and the mean nose velocity within 30cm. Note that for this analysis, we split *k* into separate parameters for casting (*k = k*_*c*_ in eq n7) and velocity (*k = k*_*v*_ in eq n9). Likewise, we labeled and varied the power parameters (n_c_ = 1 in eq n7) and (n_v_ = 4 in eq n9) from 0 - 5. We also added a coefficient in front of the *tanh* binaral term in eq n5 (k_binaral2_), and where *k*_*binaral*_ in eq n5 is relabeled as *k*_*binaral1*_. All parameters and both dependent variables significantly impacted success rate, except for the parameters *τ, σ*_*min*_, and *k*_*c*_.

To more rigorously test the effects of different levels of noise, the noise level *k*_*n*_ in eq n11 (also see step 2 in Figure 5B) was varied from 0 to 1 and the Gaussian smoothing (4mm in reported CSM variant; step 4 in Figure 5B) was varied from 0mm to 10mm. Success rate rapidly dropped off as *k*_*n*_ approached 1 or smoothing approached 0mm (Figure 7B).

To more rigorously test the effects of the 2-parameter binaral term, we re-ran the 6 reduced model variants 100,000 times each with a coefficient of *k*_*binaral2*_ =3 in front of the *tanh* term in eq n5. Again, we chose 1000 groups with 10,000 samples in each group, selected without replacement. This resulted in smaller effects of removing (-V) and (-C), but a larger effect of removing (-B), suggesting that strong binaral-sniffing may be able to compensate for removal of variable velocity and casting (Figure 7C).

In summary, reduced models revealed that individual components of the overall strategy worked together to maximize task performance, with concentration-dependent velocity and binaral-sniffing having the greatest impact on successful task performance.

## DISCUSSION

Here, we described the navigation strategies of mice during an open field odor-based spot-finding task. Over the course of 7-10 trials, mice could be trained to perform this task on a 45” × 36” open field arena. Performance of mice exceeded that of shuffled controls, indicating that mice were using odor cues to localize the odor target. Olfactory-guided behavior changed as a function of distance; specifically, 1) nose velocity decreased close to the source, 2) casting increased close to the source, and, 3) occupancy increased close to the source. Mice demonstrate reliable spot-finding within 10-45cm of the odor source. The Concentration Sensitive Model (CSM) was built to capture these observations by incorporating concentration-dependent velocity and casting terms, as well as a postulated binaral comparison term. Removing components of the CSM demonstrated that each had a significant impact on performance.

### There is a potential speed-accuracy trade-off in search strategy close to the odor spot

As mice approach an odor source, they slow down and increase their exploration time in a smaller annulus around the spot. This is similar to observations in fruit flies following a plume upwind to an odor source (Budick & Dickinson, 2006). We suggest this phenomenon may be a type of speed-accuracy trade-off (Heitz, 2014) in olfactory search behavior. Specifically, when concentration reaches some perceptual threshold, mice may “switch” into a local search strategy designed to increase accuracy at the expense of speed to target. In this local search strategy, the mouse spends more time in regions of high concentration where it is more likely to hit its target. In agreement with this hypothesis, elimination of concentration-dependent velocity decrease led to less success in reduced models (Figure 7A). An alternative explanation is that as the mouse approaches the odor source, it may reduce its running speed to avoid ‘overshooting’ the odor source.

### Binaral-sniffing facilitates search near the odor source

Use of binaral-cues for navigation has been observed in moles (Catania, 2013), snakes (Schwenk, 1994), and insects (Martin, 1965; Steck et al., 2010) suggesting it may be an evolutionarily conserved strategy. In mammals, neurons in the anterior olfactory nucleus may play a key role in integration of bilateral stimuli that are key to binaral-sniffing (Kikuta et al., 2010) as this is one of the earliest locations in which information from both left and right naris can be integrated. We suggest that binaral-sniffing cues may play a pivotal role in fine discrimination of odor identity (Uchida & Mainen, 2003), as opposed to gross determination of odor source localization in space. Once a trail’s direction, or the rough direction to a spot is determined, binaral-sniffing cues may be useful for determining the direction to cast in order to continue following the direction of the trail (Khan et al., 2012), or steer towards the odor spot. Binaral-sniffing allowed the CSM to preferentially cast in the direction of the odor source only when the odor gradient was large (i.e., near the source). Removal of binaral-sniffing led to a statistically significant decrease in CSM success (Figure 7C), an observation compatible with the model developed by Catania, 2013, where binaral-sniffing becomes a relevant strategy on steep concentration gradients. Depending on the strength of the *k*_*binaral*_ term (Eq n5), binaral-sniffing may help bias increased lateral search towards the direction of the source, allowing the nose to be “pulled” toward the source in relatively smooth arc-like trajectories.

Future research should examine if the physical distance between nostrils is correlated with the ‘effective’ nares separation of air drawn into the nasal cavity, or if the effective nares separation is determined by physical parameters such as inhalation volume, flux, and gradient. For example, inhalation in rats is largely from the lateral direction, rather than the front of the animal (Wilson & Sullivan, 1999). Pressure differences generated during sniffing inhalation at different magnitudes and frequencies (slow deep breath versus rapid sniffing) may also draw in relatively different concentrations of molecules from different distances away from the nares.

### Odor source localization may rely on cognitive mapping

Navigation to an odor source with serial-sniffing is probably intimately tied to navigation through space using the cognitive map (Jacobs, 2012). Specifically, the distance traveled between the first to the second sniff in the concentration comparison helps determine an updated heading towards the source or along the trail. In support of this view, hippocampal place cells may change their firing patterns (called remapping) to match the relocation of an orienting odor source in a circular arena (Zhang & Manahan-Vaughan, 2015). Thus, serial-sniffing is potentially more relevant to gross odor localization than binaral-sniffing. For example, to determine trail direction using binaral-sniffing most efficiently, an animal would have to be oriented roughly perpendicular to the trail so that its nares are sensing maximally different concentrations. In contrast, studies with dogs repeatedly find that serial-sniffing of 2-5 footprints is required to determine a trail’s direction (Hepper & Wells, 2005; Thesen et al., 1993). Similarly, initial odor source seeking may utilize serial-sniffing to quickly map larger spatial areas, including eliminating subspaces where no odor is detected (Vergassola et al., 2007).

### Casting is related to olfactory-guided navigation to a discrete, static odor source

Casting has been observed at frequencies of ~4 Hz in moths flying upwind during pheromone-tracking (Charlton et al., 1993), and similar head-scanning behaviors have been observed in *Drosophila* larvae, rats, moles, and humans (Baker et al., 2018). In moths, casting has been reported to occur *outside* of an odor plume when a moth is attempting to re-acquire the odor (Kennedy, 1983) and is characterized by loop-like flight trajectories (Kuenen & Carde, 1994). In contrast, here we find that mouse casting amplitude increases during approach to the odor source. The moth data suggest that casting increases whenever there are intermittent gaps in the odor signal, as they cast to re-acquire the odor plume. If this were true for mice performing this task, we would expect casting to be higher farther away from the odor source where the signal is sparse. The nature of this task, where mice seek a discrete odor object on a near-bed surface rather than an odor plume in the air, may explain these differences in behavior. While we did not find that addition of concentration-dependent casting improves performance (Figure 5B), we did not prove that this was true for all parameter sets in the CSM model. Therefore, a set of untested parameters may explain the lack of improvement with addition of concentration-dependent casting.

### Complexities to address in future studies include decision-rules and individual variability in strategy utilization

Our data suggest that mice may use multiple search strategies during an open field odor-based spot-finding task depending on their distance from the source. We speculate that the relative contribution of strategies may also be variable between individuals, as has been observed during fruit fly navigation during odor presentation (Tao et al., 2019). This may mean a specific instantiation of our model (with one set of parameters) captures behavior for one individual mouse, but fails to capture behavior for some other subset of mice. For example, we found that the *k* parameter could be used to ‘tune’ aspects of the model’s trajectories which may better fit the data for a subpopulation of mice. A much larger cohort of animals than was used here would be necessary to evaluate this idea; however, the general principle is that similar-looking ethologically-relevant behavior can emerge from different model permutations.

A limitation of the current model is that it incorporates an ‘all or nothing’ decision at each sniff. It may be more appropriate to use a probabilistic decision rule to combine binaral- and serial-sniffing into a unified output. Also, a model where pausing emerges during epochs of uncertainty in sampling of the odor concentration gradient may be advantageous. Additionally, the model is currently limited to detection of one odorant—it would be useful to distinguish different features of diverse odorants. Finally, stimulus models with ‘zeros’ distributed throughout the simulated odor landscape produced high failure rates. Developing a model that is robust to this type of intermittency would be a logical next step.

We show that the CSM is capable of using serial concentration sampling of gradients to locate odors, even when the odor landscape has many local minima and maxima surrounding the odor source (e.g., Figure 5B, step 4). While the CSM is capable of finding the odor source when the odor is consistently above threshold, it does not attempt to capture mouse behavior in the environmental regime where the odor concentration is below the animal’s detection threshold. Rather than attempting to model mouse behavior in this subthreshold condition, the CSM engages in a correlated random walk that may bring it in contact with suprathreshold odor concentrations. The mouse may be engaging in more complex navigation behavior in this dilute regime, including mapping of the odor environment. Models that attempt to navigate complex odor landscapes without gradient ascent have their own drawbacks. The model of Balkovsky and Shairaman, 2002 requires an additional wind cue - anemotaxis - to function properly. The model of Vergasolla, *et* al, 2007 builds a cognitive map of odor source location likelihood; it utilizes both sparse odor detection events and frequent subthreshold observations to update its map of source location likelihood.

In summary, we discover that mice vary their search strategy, including casting, during active navigation toward a discrete odor object, modulating features of behavior as a function of distance from the target. A concentration-sensitive model successfully reproduces mouse behavior, while allowing for the analysis of the relative contributions of binaral-sniffing, concentration-dependent velocity, and concentration-dependent casting amplitude. Future work is needed to experimentally dissect how these variables impact mouse navigation and whether additional parameters are important in successful navigation to a static odor source.

## ACKNOWLEDGEMENTS

We thank Gregory LaRocca for technical help, and members of the Urban laboratory as well as Yu Chen, Robert Kass, Nour Riman, Erin Connor, and John Crimaldi for helpful discussions. We thank Kathy Nagel and Jonathan Victor for helpful comments on an earlier draft of the manuscript. A special thanks to Peter Jones for designing and building the behavior table used in this experiment. Thanks to Chris D. Rand from Aurora Scientific for technical help with the PID.

## SUPPLEMENTAL CAPTIONS

**Figure S1.**
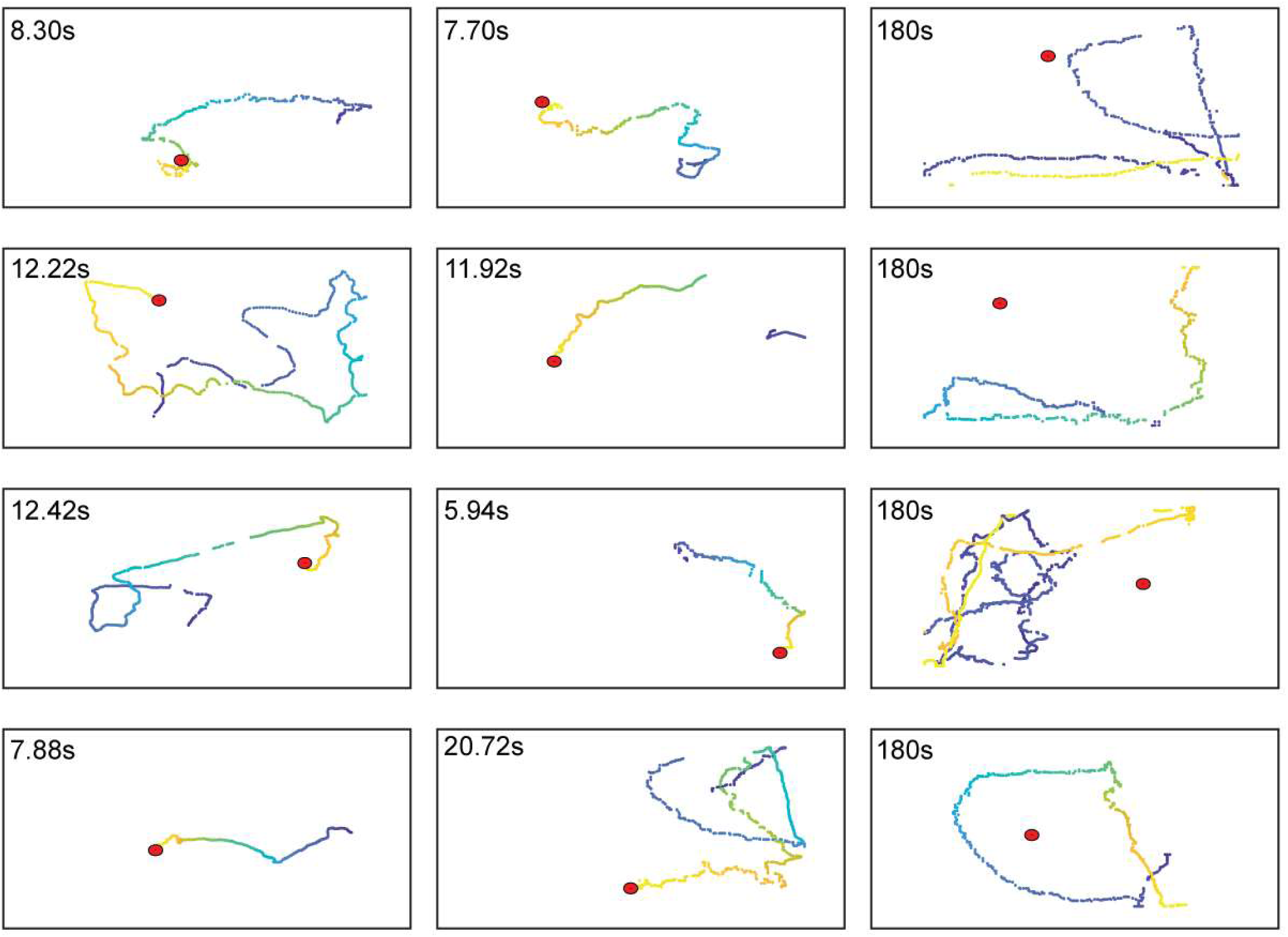
Example Traces. Successful trials (left and center panels) have the time to spot indicated by the text. The relative time of each position sample is indicated by the color. Unsuccessful trials (right panel) have the total trial time indicated by the text. Color represents the relative time of each position sample. Spot positions are indicated by red circles.

**Figure S2.**
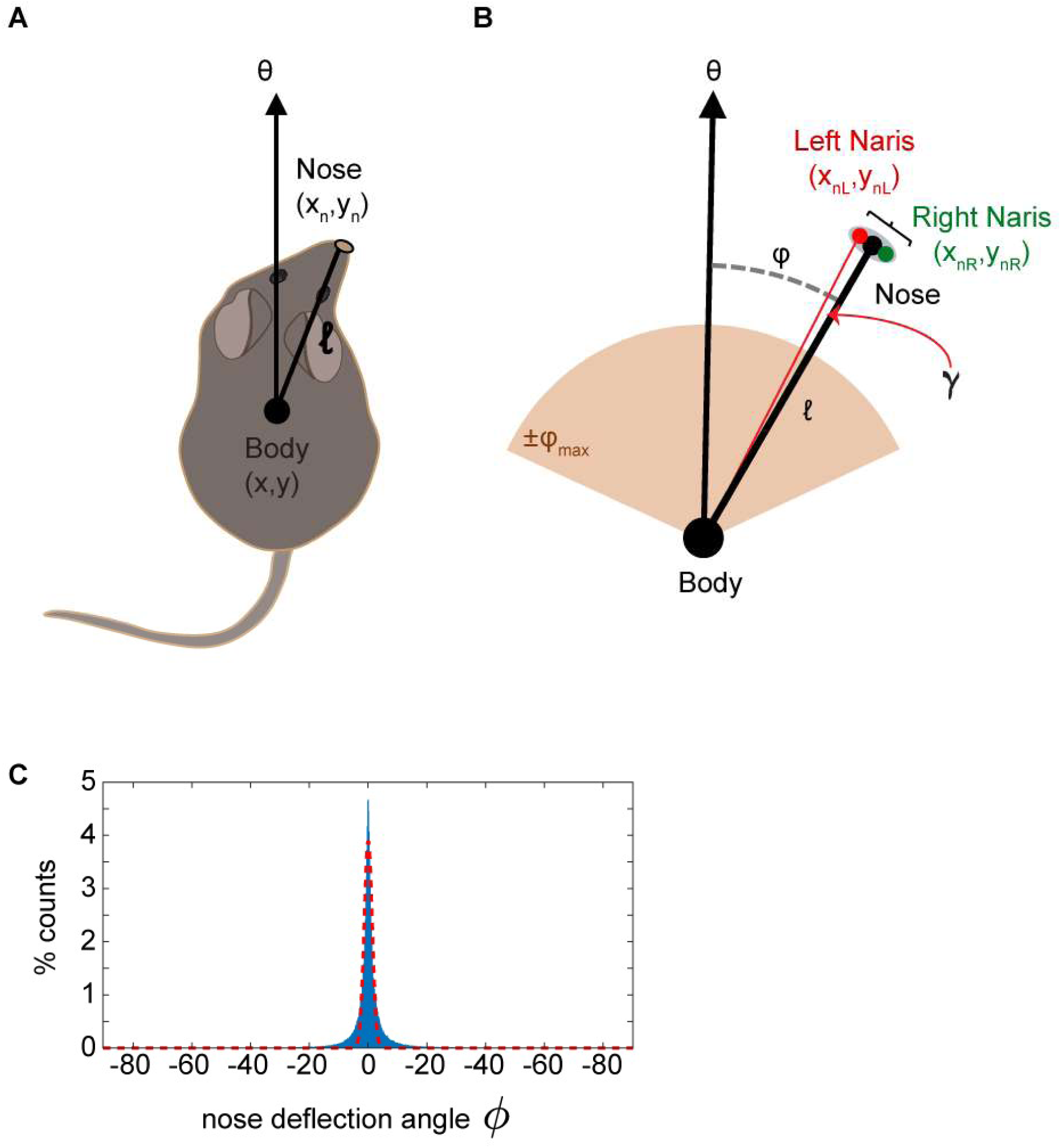
Detailed model geometry. **A)** Model specification of the body position (x,y), and the body angle *θ* superimposed on a diagram of the mouse. The nose position (x_n_,y_n_) is the mean position between the left and right nares. **B)** Nose angle *ϕ* (gray dashed line) is the angle between *θ* and nose position. Nose angle *ϕ* is constrained by a maximum angle *ϕ*_*max*_. The head-body length *ℓ* is the distance between the body position and the nose position. Nose position is sub-divided into two nostrils (left indicated by red, right indicated by green), separated by a distance *d*_*nares*_. The angle from the center of the nose to the edge of the nostril is γ. **C)** The distribution of mouse nose deflection angles *ϕ* (blue bars) is approximately Gaussian (dashed red line; a*exp(-((x-b)/c)^2); a=0.04, b=0.06, c=1.92).

**Figure S3.**
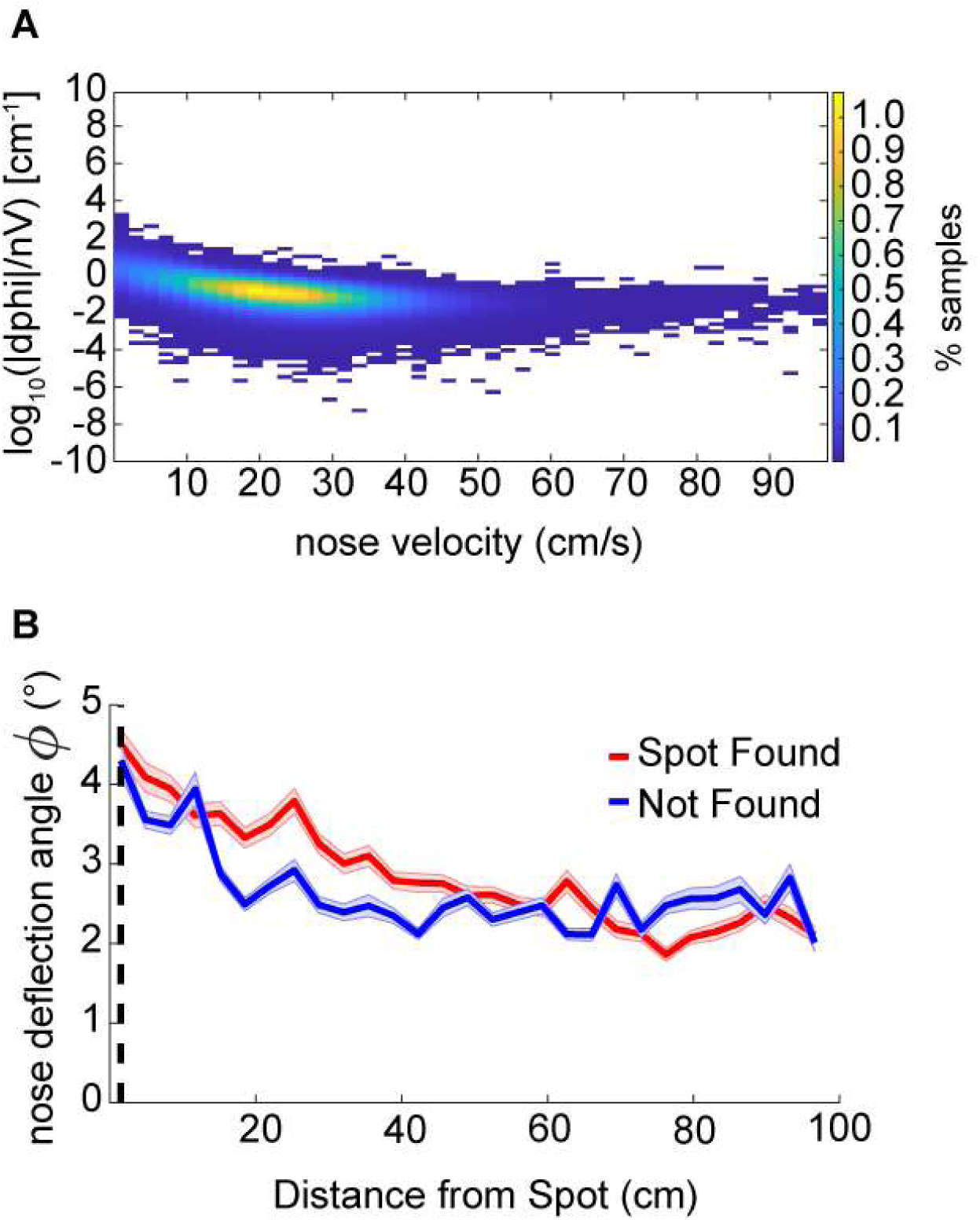
Additional Casting Analyses. **A)** Curvature as measured by log_10_(|dPhi|/nV) depends on nose velocity. The colormap indicates the percentage of samples in each bin. **B)** Averaged across successful trials, nose deflection angle *ϕ* increases as the distance to the spot decreases (line=mean±s.e.m; ANOVA; p=2.18E-56, F=12.82, df=28).

**Table S1.**
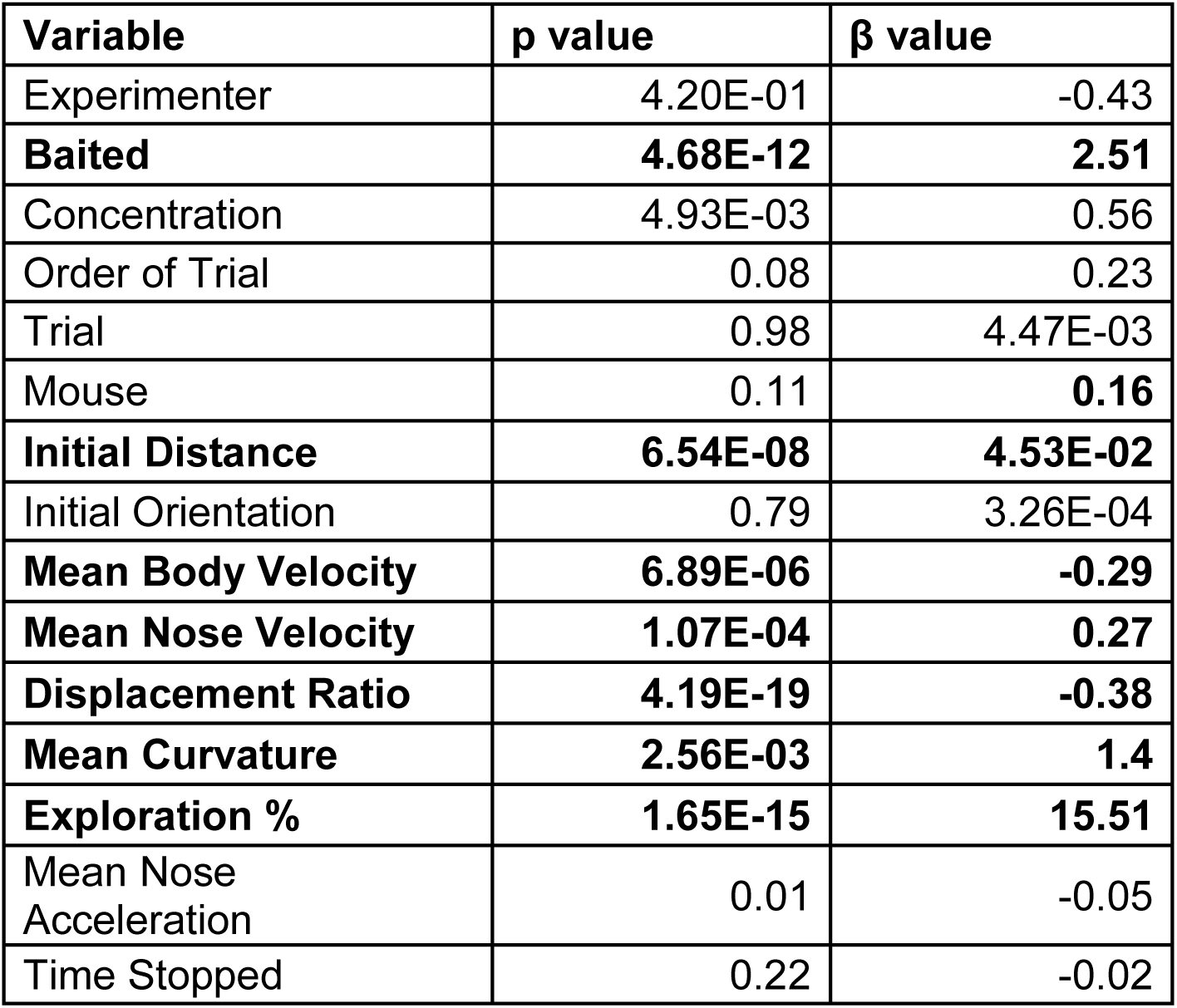
Logistic regression analysis on spot-finding success in mice. Significant variables in bold.

**Table S2.** Logistic regression analysis of CSM parameters on spot-finding success. Significant parameters are in bold.

| **Variable** | **p value** | **$\beta$ value** | **Range** |
| --- | --- | --- | --- |
| **d<sub>nares</sub>** | **1.00E-50** | **0.54** | **0.01 - 1cm** |
| **k<sub>c</sub>** | 4.84E-03 | -0.1 | 0 - 1 |
| **k<sub>binaral1</sub>** | **6.86E-26** | **1.27E-03** | **0.01 - 300** |
| **k<sub>binaral2</sub>** | **7.33E-106** | **0.16** | **0 - 5** |
| **k<sub>v</sub>** | **2.63E-47** | **0.62** | **0 - 1** |
| **$\ell$ (neck length)** | **1.35E-191** | **0.26** | **0.01 - 10cm** |
| **Mean Curvature (dist < 30cm)** | **6.13E-09** | **0.6** | **Dependent Variable** |
| **Mean Nose Velocity (dist < 30cm)** | **2.30E-19** | **-0.016** | **Dependent Variable** |
| **n<sub>c</sub>** | **4.67E-09** | **-0.043** | **0 - 5** |
| **n<sub>v</sub>** | **4.76E-160** | **0.22** | **0 - 5** |
| **$\sigma_{\max}$** | **1.11E-39** | **0.77** | **0.01 - 1** |
| $\sigma_{\min}$ | 2.50E-01 | 0.074 | 0.01 - 1 |
| $\tau$ | 6.40E-01 | -0.02 | 0.01 - 1 |
| **V<sub>max</sub>** | **9.26E-54** | **0.03** | **20 - 40cm/s** |

## ABBREVIATIONS

CSM: concentration-sensitive model
CRW: correlated random walk
OU: Ornstein-Uhlenbeck
Δt: change in time (s)
ℓ: nose-to-body length parameter (cm)
*v*_*max*_: maximum velocity parameter (cm/s)
k: concentration-dependent parameter
σ_min_: concentration-dependent parameter
σ_max_: concentration-dependent parameter
k_binaral_: binaral-olfaction parameter
d_nares_: inter-nostril parameter (cm)
ANOVA: Analysis Of VAriance
LED: light emitting diode
|dPhi|/nV: curvature
x’: derivative (of x)
x_n_: nose coordinate
θ: heading of model body (degrees)
φ: heading of model nose relative to body (degrees)
C_t_: concentration magnitude (A.U.)
η_t_: multiplicative uniform noise
(-C): CSM model without concentration-dependent casting, *σ* = *σ*_*min*_
(-B): CSM model without binaral-sniffing component, *k*_*binaral*_=0
(-V): CSM model without concentration-dependent velocity, *v*=*v*_*max*_
IR: infrared

